# Courtship song differs between African and European populations of *Drosophila melanogaster* and involves a strong effect locus

**DOI:** 10.1101/2024.05.14.594231

**Authors:** Matthew J. Lollar, Elizabeth Kim, David L. Stern, John E. Pool

## Abstract

The courtship song of *Drosophila melanogaster* has long served as an excellent model system for studies of animal communication and differences in courtship song have been demonstrated among populations and between species. Here, we report that flies of African and European origin, which diverged approximately 13,000 years ago, show significant genetic differentiation in the use of slow versus fast pulse song. Using a combination of quantitative trait mapping and population genetic analysis we detected a single strong QTL underlying this trait and we identified candidate genes that may contribute to the evolution of this trait. Song trait variation between parental strains of our recombinant inbred panel enabled detection of genomic intervals associated with six additional song traits, some of which include known courtship-related genes. These findings improve the prospects for further genetic insights into the evolution of reproductive behavior and the biology underlying courtship song.

## INTRODUCTION

Communication, the process of information transfer through signal production and recognition, is abundant across the animal kingdom. There are many forms of communication in nature, from chemical interactions between bacteria (Keller & Surette 2006), to intricate mating displays of suboscine passerines (Prum 1990), or the complexity of human language (Lieberman 1971). Accompanying this diversity of form is a diversity of purpose: communication is used by animals in mate choice and conspecific recognition (Anholt *et al*. 2020), antagonistic displays (Maynard Smith 1979), and cooperative behavior such as territory defense and foraging (Fitzgerald & Peterson 1988). Often, the proper transmission of information from signaler to receiver has important consequences for the fitness and reproductive success of an individual (Maynard Smith 1979; Anholt *et al*. 2020), underpinning its importance in both standing diversity and the evolutionary history of populations. Acoustic communication has evolved independently over 20 times across insects (Greenfield 2016) and is an important component of reproductive behavior (Leonhardt *et al*. 2016).

Male *Drosophila melanogaster* “sing” to females by vibrating their wings, typically as unilateral extension of a single outstretched wing (Ewing & Bennet-Clark 1968). In *D. melanogaster*, song consists of two primary modes: a series of continuous sinusoidal waves or hums (sine song) and trains of more punctuated wave clusters (pulse song) with amplitude and pitch typically exceeding those produced by sine song (Swain & Philipsborn 2021). Furthermore, two modes of pulse song have been identified based on the pitch of the pulse song: fast and slow pulse modes (Clemens *et al*. 2018). Female *Drosophila* typically become more receptive to mating upon hearing song (von Schilcher 1976) and may distinguish between subtle differences in song components. The active female responses to courtship that encourage or discourage copulation underpin the role of song in the reproductive success of male flies (Cook 1973; Riabinina *et al*. 2011; Laturney & Moehring 2012).

Male courtship song has been characterized in many *Drosophila* species, (O’Grady & DeSalle 2018), providing deep historical knowledge in a system benefiting from excellent genetic tractability (St. Johnston 2002; *Drosophila* 12 Genomes Consortium 2007). The acoustic properties of pulse song, such as the interpulse interval (IPI) typically receive the greatest attention (Ritchie *et al*. 1994; Talyn & Dowse 2004; Stern 2014), although components of sine song also vary among closely related species (Ding *et al*. 2016). Pulse song is highly variable across species and has been implicated in mate and conspecific recognition (Spieth 1952; Bennet-Clark & Ewing 1969; Watanabe & Kawanishi 1979; Cowling & Burnet 1981; Talyn & Dowse 2004; Ding *et al*. 2019), although song does not appear to contribute to mate choice in every species (Richie *et al*. 1999; Iglesias & Hasson 2017).

The genetic architectures and underpinnings of variation in courtship song and other premating isolating mechanisms are poorly understood. Early studies supported the view that variation in these behaviors was caused by a small number of loci of large effect (Mackay 2001; Mackay *et al*. 2005). Arbuthnott (2009) found, for instance, that among 36 studies in insects (predominantly *Drosophila)* on courtship traits contributing to premating isolating barriers, 25 studies considered traits to be influenced by a few loci of large effect. However, quantitative trait locus (QTL) studies are likely biased toward detecting large effect loci (Rockman 2012). More recent studies have revealed that the genetic architecture of premating barriers is likely more complex (Mackay 2009; Arbuthnott 2009; Oh *et al*. 2012; Shahandeh & Turner 2020). The number of loci contributing to variation in a trait involved in premating isolation can also have implications for the rate of speciation between species (Gavrilets & Vose 2007), providing additional incentive to study the evolution of such phenotypes.

Differences between well-isolated species in reproductive behavior may or may not have contributed to the initial reproductive isolation of those species (Price 2002). On the other hand, within-population variation in such traits may be distinct in its genetic architecture and basis from evolutionary trait changes (Arbuthnott 2009). Whereas, evolved differences in courtship traits between interfertile taxa, including recently diverged populations of the same species, may hold particular interest for the study of reproductive isolation.

*D. melanogaster* originated in sub-Saharan Africa and has spread nearly worldwide (Lachaise & Silvain 2004, Lachaise *et al*. 1988, Keller 2007). The demographic history of the species is well-studied (e.g. Sprengelmeyer *et al*. 2020) and striking genetic and phenotypic changes have occurred on a relatively short evolutionary time-scale. Genetic differentiation is in part due to at least one moderate population bottleneck during the initial migration out of sub-Saharan Africa (Begun & Aquadro 1993, Sprengelmeyer *et al*. 2020), as well as locally adaptive changes in response to novel environmental pressures (David & Capy 1988, Fabian *et al*. 2015; Lack *et al*. 2016a). Genes under selection as a consequence of adaptation often have neuronal or morphological and developmental functions (Fabian *et al*. 2015; Lack *et al*. 2016a; Pool *et al*. 2017). Such changes could have either specific or pleiotropic effects on courtship traits such as song, in light of the multifaceted roles of many genes involved in courtship behavior (Gleason 2005; Greenspan & Ferveur 2000).

While some components of courtship song variation within *D. melanogaster* have been studied (Colegrave *et al*. 1999; Gleason *et al*. 2002), there have been no prior comprehensive intraspecific studies of many courtship song components. Here, we utilize panels of naturally-derived *D. melanogaster* from two geographic origins, Zambia and France, and high-throughput methods of recording and analyzing courtship song (Arthur *et al*. 2013; Arthur *et al*. 2021) to study the genetic basis of song differences between populations. We find that the proportion of slow and fast pulses (“pulse mode preference”) differs significantly between populations. We use a panel of recombinant inbred lines (RILs) derived from reciprocal crosses between France and Zambia strains to map pulse mode preference as a quantitative trait, as well as mapping eight additional song traits that differ significantly between RIL parental strains. Finally we complement our mapping analysis with population genetic screens across QTL intervals, highlighting the most genetically differentiated loci between populations to identify additional genes that may underline song variation and evolution. Collectively, our results reveal that naturally evolved differences in male song components exist between populations within the *D. melanogaster* species and include moderately strong effect loci, opening the door to targeted genetic and neurophysiological studies of their molecular basis.

## MATERIALS AND METHODS

### Drosophila husbandry and stocks

Two distinct types of strains were used in this study. Strains used to measure trait differences between population samples were (8 generation) inbred lines from wild-derived isofemale stocks, collected at locations described in Pool *et al*. (2012). We used a recombinant inbred line (RIL) panel generated in the lab and derived from a reciprocal cross between two inbred strains, one each from our France (FR) and Zambia (ZI) collections (FR320N and ZI418N), for QTL mapping. Following 12 generations of intercrossing, isofemale lines were established from these interbred flies and inbred for 5 generations to establish the final panel, from which 305 recombinant strains were included in the present study. Females used in mating assays were derived from a single France strain, FR54N. We chose this strain because European females show less preference across populations compared to some southern African strains (Grillet *et al*. 2012; Hollocher & Wu, 1996; Wu *et al*. 1995), who tend to disfavor non-African males, and female response can influence the expression of song (Riabinina *et al*. 2011; Laturney & Moehring 2012).

Flies were maintained as stocks in laboratory conditions on food prepared in batches of 4.5 L water, 500 mL cornmeal, 500 mL molasses, 200 mL yeast, 54 g agar, 20 mL propionic acid, and 45 mL tegosept 10% (in 95% ethanol). Flies used in assays were collected under carbon dioxide anesthesia and aged in one of two 12 hour light/dark controlled incubators, separated in cycle by two hours to allow additional assays each day. Incubator temperatures were not strictly controlled but measured daily and ranged from 23-25C. Females were collected and aged in plastic fly food bottles containing approximately 50 females per bottle. Males were collected and aged individually in glass fly food vials. The age of collected males ranged from 0-24 hours post-eclosion and was not strictly controlled for prior matings during this time. Previous matings can affect male mating success (e.g. Sinclair *et al*. 2021), although the frequency of matings among our males may have been low due to the 0-24 hours age range of the females (Manning 1967). Whereas, prior courtship attempts may have been more variable among the collected males, given that males begin to court as early as 12 hours post-eclosion (Strömnæs & Kvelland 1962) and may court virgin females (Dukas 2005). The collection of tens of males from each fly strain (see below), and the averaging of their song trait values, should dampen individual-level variation induced by such factors.

### Song recording

Male song was recorded on a multi-channel acoustic recording chamber (Arthur *et al*. 2013), at Janelia Research Campus from June-July of 2022 (Table S1). Males used in assays were collected 0-24 hours post-eclosion, isolated into individual vials, and aged for 3-5 days. Males were recorded in the presence of a single unmated female, collected 0-6 hours post-eclosion, and aged an additional 12-24 hours before use in assays. This age was selected as an optimal range to allow aging of females past a “young nymph” stage, without allowing them to become old enough to be receptive to mating (Manning

1967), which would diminish the duration of captured song before copulation (observationally, males that initiated song typically sang throughout the entire duration of recording if they did not mount successfully). Flies were mouth aspirated into recording chambers, and assays were performed within the first two hours from lights on. Male flies were recorded for a consecutive duration of 30 minutes. Replicate assays were performed such that strains were assayed in sets of 6-8 males across 3-5 unique days, for a total target sample size of 24 for each RIL, and 50-60 for each inbred strain (Table S1).

### Song analysis and software protocols

Song data was automatically segmented and analyzed without human intervention using SongExplorer (doi:10.1101/2021.03.26.437280) to annotate acoustic components of song. A previously trained *D.melanogaster* model detected four principle sounds (sine song, pulse song, ambient noise, other), trained using audio collected on the recording instrumentation used in this study. Samples were recorded at a sampling rate twice that of the provided model, so raw audio files were downsampled at a rate of one half using the “signal” function in the python open-source package SciPy (version 1.13) for proper segmentation (Virtanen *et al*. 2020).

Segmented song parameters were quantified using BatchSongAnalysis (https://github.com/dstern/BatchSongAnalysis) with small modifications for individualized trait parameterization (https://github.com/mjlollar/song_work), MATLAB version 9.13 (R2022b) (The MathWorks Inc. 2022). The numbers of song bouts, sine and pulse song durations, interpulse interval characteristics, pulse/sine carrier frequencies and pulse/sine train lengths were measured as described in Arthur *et al*. (2013). Pulse and sine song amplitude were measured by calculating the square root of the mean of the squares of all pulses in a song (output as arbitrary units). Transition probabilities were calculated using the Matlab “transProb()” function for pairwise measurements of events (The MathWorks Inc. 2022). Slow and fast pulse characterization was available thanks to previous classification training in the model provided and used methods similar to those described in Clemens *et al*. (2018) to differentiate and classify pulse modes. Interpulse intervals were estimated independently of pulse carrier frequency by fitting an envelope to each pulse to estimate pulse duration and calculating the interpulse interval as the time from the end of one pulse to the center of the next pulse. Here, pulse event carrier frequency was measured as the modal frequency detected by SongExplorer. Further discussion on the characterization of carrier frequency can be found in Ding *et al*. (2019).

Replicates with fewer than one second of song per minute, or those with an aberrantly high sine song rate per minute (recordings with >55 seconds of characterized sine song appeared to represent a failure of recording equipment) were excluded from analysis. Additionally, replicates with zero values for a trait were not considered when calculating strain mean for that trait, as a zero value from BatchSongAnalysis was typically the result of an inability to properly measure trait values (due to minimal sampling throughout the assay, etc.). Trait values for strains were represented as the mean value of the total remaining replicates.

Prior to filtering, the number of replicate inbred and RIL samples totaled 860 and 7,152, respectively. After the filtering described above was applied to raw data, the total number of inbred song recordings were 810, with an average replicate number of about 43 recordings per strain. The total number of RIL song recordings totaled 5,532 after filtering, with an average of about 19 recordings per RIL line.

### Data analysis of processed song recordings

Differences in the population mean trait values were quantified for each trait using Mann-Whitney (MW) tests using the two-sample wilcox.test() function in R (R Core Team 2023; https://github.com/mjlollar/song_work). To statistically control the testing of multiple traits, we used a permutation-based approach to correct for multiple testing that accounted for non-independence among some traits (Figure S1). 10000 permutations were performed in the following design. For each replicate, the identities of each strain were randomly shuffled and drawn without replacement to represent 10 FR strains and 7 ZI strains (matching sample sizes of the empirical data). MW tests were calculated between these groups and recorded. We then determined the false positive rate (the likelihood the empirical MW p-value occurs by chance) as the proportion of permutation replicates that gave a p-value as low or lower than the empirical value. Correlation statistics were generated in R using Pearson’s test (R Core Team 2023).

For a trait showing evidence of population differentiation from the above methodology, we also calculated Q_ST_ as a measure of quantitative trait differentiation (Lande 1992; Spitze 1993). To indicate whether this trait might fall outside the range of neutral population differentiation, we compared the estimated Q_ST_ to the distribution of F_ST_ at individual SNPs (which conservatively maximizes the variance of F_ST_), calculated from allelic sample sizes identical to the number of strains from which phenotypic data was obtained. F_ST_ distributions were calculated using available genomes from the *Drosophila* Genome Nexus (Lack *et al*. 2016a; see below).

### QTL mapping based on RIL ancestries

Genomic sequencing of the RIL panel is described in da Silva Ribeiro *et al*. (2024), which also documents the calling of parental strain ancestry. Briefly, for each RIL, homozygous and heterozygous parental strain ancestry calls were assigned using Ancestry HMM (Corbett-Detig & Nielsen 2017) on windows defined by intervals containing 2,500 SNP genotype differences between the two parental strains (which were used as the parental panels for Ancestry HMM). This window size provides an abundance of variants from which to infer ancestry in any particular window, but given that they average 5 kb in length, they are still much smaller than the typical expected length of ancestry tracts, which should tend to be roughly on the megabase scale after 12 generations of interbreeding.

QTL mapping was performed using parental strain ancestry calls as marker genotypes. We used a modified version of r/QTL (Broman *et al*. 2003) that allowed us to use an additive QTL model, rather than allowing each genotypic class to have an independent mean trait value (da Silva Ribeiro *et al*. 2024), in light of the smaller counts of some ancestry genotypes observed in many genomic windows. The primary model tests the alternative hypothesis of a single QTL against the null hypothesis of no QTL, while using phenotypic permutations (n=10,000) to control for genome-wide multiple testing. More explicit details of the pipeline can be found in da Silva Ribeiro *et al*. (2024), as well as source code at https://github.com/ribeirots/RILepi. We note that this permutation approach, along with the averaging of multiple individual trait values per RIL, should minimize the potential influence of skewed trait distributions on our QTL mapping.

The above analysis yields a genomic LOD score landscape for any given song trait. However, when this landscape is complex or ragged in its topography, in practice it can be difficult to assess which local peaks to consider as distinct QTLs. We therefore implemented a series of criteria to determine which local peaks to integrate into the same QTL region, and also to denote which regions within a broad/complex QTL may have the greatest likelihood of containing a causative locus. Preliminary QTL peak regions were defined based on contiguous runs of windows above the genome-wide LOD significance threshold. A pair of initially separate peaks separated by a valley with a minimum LOD value of greater than the shorter peak’s maximum LOD score minus 1.5 were merged into the same peak region – since a simple confidence interval (CI) for the shorter peak, based on a 1.5 LOD drop, would end up including the taller peak. After defining and integrating peaks in this manner, we accounted for peaks with complex LOD landscapes by differentiating between “inclusive” and “restrictive” CIs. Within a contiguous run of windows above the LOD significance threshold, the inclusive CI was defined as the full span encompassing all local windows in which the local LOD score was greater than the region’s maximum LOD score minus 1.5. A QTL region’s inclusive CI was therefore contiguous by definition, and could encompass some windows with LOD scores less than the maximum LOD minus 1.5.

In contrast, restrictive Cis were defined as only the subset of windows within the inclusive CI that had LOD scores greater than the maximum LOD minus 1.5 (https://github.com/mjlollar/song_work). A QTL region’s restrictive CI could therefore encompass multiple non-contiguous runs of windows representing different LOD sub-peaks within the same QTL region as defined above. Based on the LOD scores of their constituent windows, restrictive Cis were considered to be the most likely intervals to contain a causative locus, and were therefore the focus of downstream analysis as described below. Windows yielding a RIL panel-wide ancestry proportion above 90% for either parental strain were excluded from the mapping analysis, and confidence intervals (Cis) encountering such regions were extended until encountering an included window that ended the CI based on the criteria detailed above.

### Residual Mapping of Correlated Traits

In cases where we separately performed QTL mapping on two correlated traits, we investigated whether overlapping QTL might reflect pleiotropic effects of a single locus. We performed a limited set of residual trait mapping analyses using linear regression to find each RIL’s residual trait values for the second trait after adjusting for its observed value for the first trait (here, treating the trait with the greater QTL LOD score as the first trait). QTLs and confidence intervals were identified using identical methods described in the previous sections. Linear regression was performed in python using the SciPy *linregress* package.

### Population Genetic and Candidate Gene Analysis of QTL regions

Previous population genomic data available from the *Drosophila* Genome Nexus (Lack *et al*. 2016a) was used to generate a population genetics database. We used all available Zambia and France genomes for a total of 197 and 96 strains, respectively. Under the hypothesis that the two population-differentiated song traits may have been influenced by population-specific selection, we assessed three window-based population genetic statistics across the restrictive Cis of each QTL region for those traits: 1) *F_ST_FullWin,_* which represents the estimated *F_ST_* value of a window using all variable sites within the window; 2) *F_ST_MaxSNP_*, the maximum *F_ST_* value of any variable site within a window (da Silva Ribeiro *et al*. 2022); and 3) Comparative Haplotype Identity (*χ*_MD_), which tests for evidence of local adaptation by comparing the summed lengths of pairwise identical haplotypes within one population versus another (Lange & Pool 2016).

We then identified “candidate genes” within QTL Cis (using the restrictive CI definition above, including only the windows within an extended CI that had LOD scores greater than the peak value minus 1.5). We associated genes with windows if any of their exons (coding or non-coding) overlapped a given window, or if an exon of this gene was the first exon to the left or right of this window’s boundaries (in order to account for potential regulatory regions of a given gene). We included as candidates any genes that had been previously implicated in courtship behavior. A list of 92 such genes was generated using genes sourced from the Gene Ontology (GO) database category “male mating behavior” (ID GO:0060179) as well as additional genes identified from cursory literature searches (Table S2).

For traits with evidence of population differentiation, we additionally included as candidates genes with a documented role in wing development or the nervous system. Wing genes included the GO categories “wing disc development” (GO:0035220) and “wing disc morphogenesis” (GO:0007472), for a total of 743 genes. Neuronal genes included the GO categories “nervous system development” (GO:0007399) and “nervous system process” (GO:0050877), for a total of 1,810 genes. For these same differentiated traits, we also annotated genes with evidence for local adaptation, in that they were associated with at least one window that fell within the top 1% quantile of values for either *F_ST_FullWin_, F_ST_MaxSNP_*, or *X_MD_*. A gene was considered to be associated with an outlier window if at least one of its coding exons overlapped the window boundaries or represented the nearest neighboring exon (in either direction).

## RESULTS

### AFRICAN AND EUROPEAN POPULATIONS OF D. MELANOGASTER MAY DIFFER IN MULTIPLE COMPONENTS OF PULSE SONG

We first sought to determine whether any quantitative aspects of male courtship song varied between two *D. melanogaster* populations: one from within the species’ inferred ancestral range (Siavonga, Zambia) and one from Europe (Lyon, France). We divided components of male song into a range of acoustic properties, each considered as a single quantitative trait. To first identify population-level differences in specific song traits, we selected 10 France-derived and 7 Zambia-derived inversion-free inbred strains from previous collection efforts (Pool *et al*. 2012), and recorded male songs in 30 minute intervals using 40-60 males per strain (raw individual recording measurements available in Tables S3 and S4). Strains were assigned single trait values for each song behavior representing the mean of individual replicates (Table S5).

In total, 37 traits were analyzed, many of which were found to have significant positive or negative correlation, often between traits with *a priori* expectations of possible correlation (e.g. the amount of song per minute and the number of bouts per minute are positively correlated) (Figures 1, S1, S2). Among traits not correlated by definition, we noted the strongest positive correlation between the sine and pulse amplitude traits, which could suggest pleiotropy for song amplitude traits. Additionally, there was a strong negative correlation between pulse carrier frequency and pulse mode preference, which could be explained by a difference in amplitude between the two types of pulse song, which has previously been demonstrated to correlate to carrier frequency (Clemens *et al*. 2018). Interestingly, the FR and ZI cohorts of inbred song had slight variation in which traits were correlated (Figure 1A).

**Figure 1:**
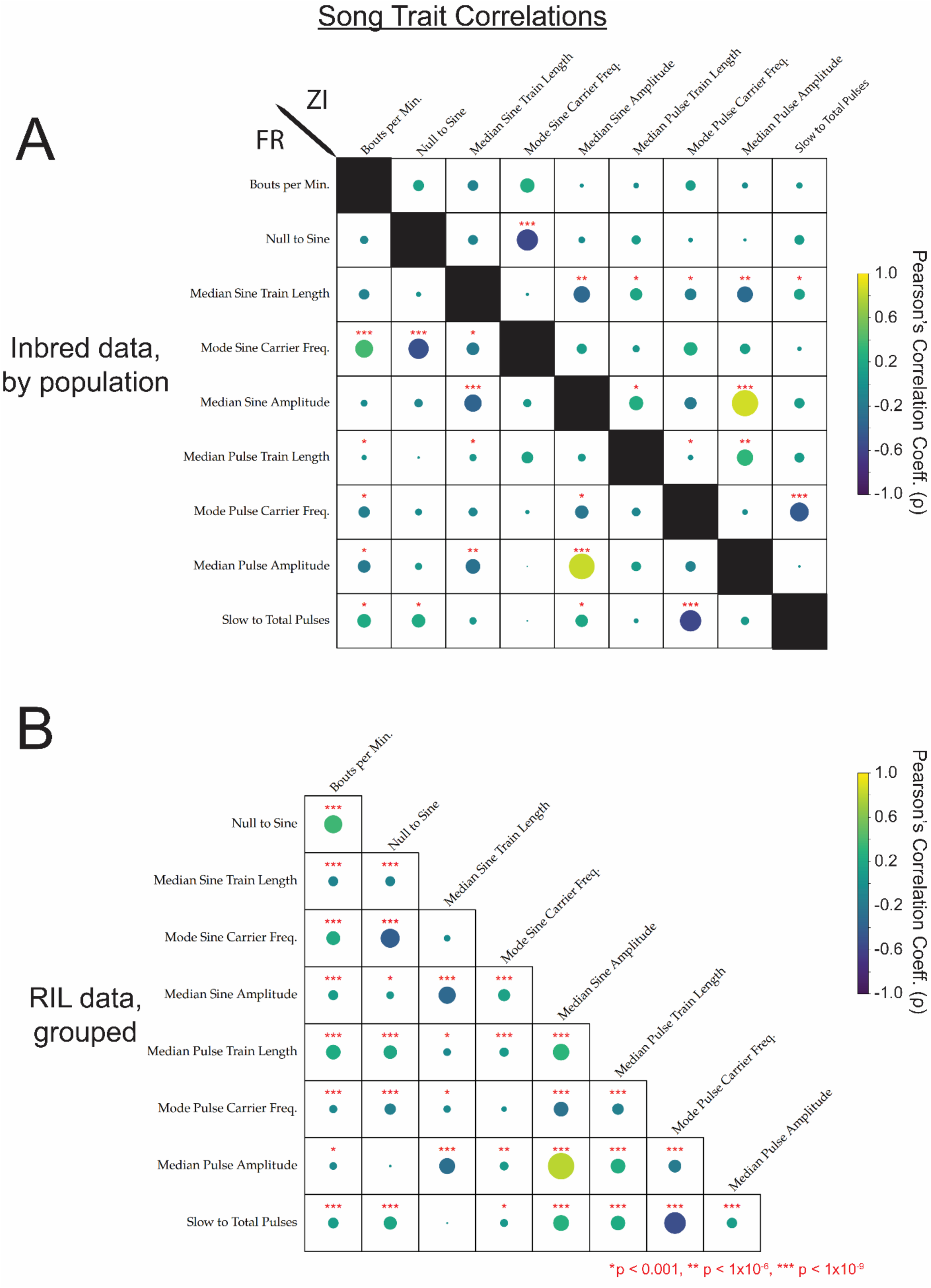
Correlation analysis between traits mapped from male song recordings reveals several positively and negatively correlated traits. **A.** Correlations among inbred strains, divided by population (France bottom-left, Zambia top-right). **B.** Correlations among RILs. In all data sets, the strongest positive correlation was between pulse and sine amplitude traits. Circle size indicates the relative strength of significance in Pearson’s (r) covariance between traits, where the presence of a circle indicates minimally a p-value of p < 0.05, and correlations with an estimated multiple-testing corrected significant p-values of p < 1.0 × 10^−3^, p < 1.0 × 10^−6^, and p < 1.0 × 10^−9^ are denoted with asterisks (*, **, and ***, respectively).

Population mean trait values were compared statistically between France and Zambia strain panels to test for significant trait divergence between populations in each examined song characteristic (Figure 2; Table S5). Two related traits, described by Clemens *et al*. (2020), had significant raw Mann-Whitney (MW) p-values for the population comparison: the number of “fast” pulses (p = 0.0185) and fraction of “slow” pulses to total pulses (p = 0.0068; Figure 2A; Table S5). Hereafter, we focus on the latter of these correlated traits (Figure 1AB) in light of its clearer signal of differentiation. Aside from these two characteristics of pulse type, the only trait with even a marginal raw p-value (p = 0.087) was the median amplitude of all pulses (Figure 2B; Table S5); France strains were associated with somewhat higher (louder) pulse amplitudes than Zambia, on average.

**Figure 2:**
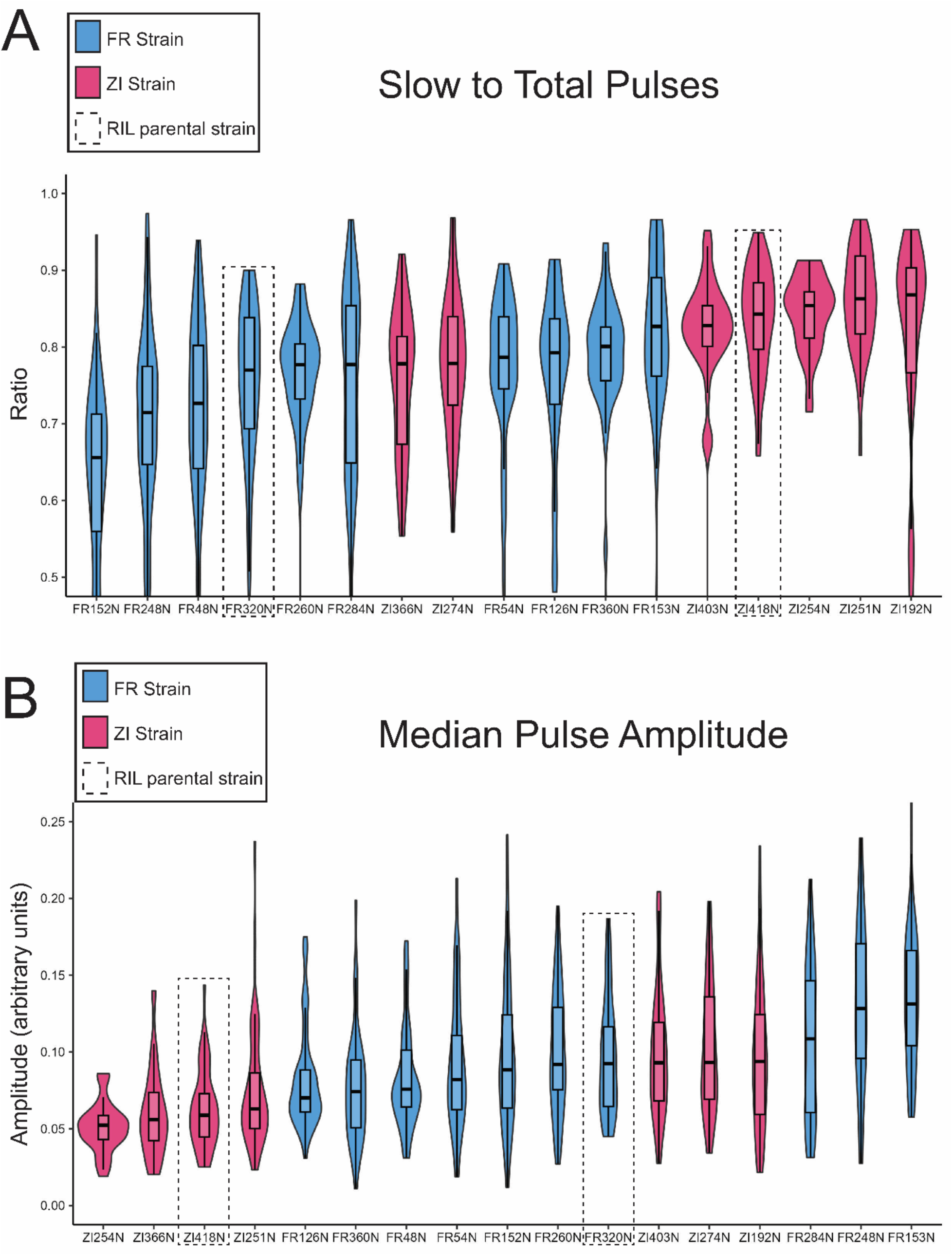
Zambia and France D. melanogaster males display differences in pulse mode preference (slow to total pulses) and in pulse amplitude. Violin and boxplots for France (Blue) and Zambia (Red) males are displayed, with the middle line in boxplots representing median values, and inbred strains ordered by those median values. Parental strains used in the establishment of the RIL panel are highlighted by dotted boxes. The pairwise differences in both traits between parental strains is of similar magnitude and direction to that of the population trend, suggesting that some of the genetic variation contributing to phenotypic differences between populations may be represented among the RIL panel.

We applied a permutation-based multiple test correction to MW p-values (Table S5). Here, we found that the ratio of “slow” pulses to total pulses had the lowest least likelihood of false positive discovery (∼10%) and represented the greatest difference in population trait differentiation (Figure 2; Table S5). Thus, the rate of slow to total pulses is likely to represent a genuinely differentiated trait between the Zambia and France populations. In contrast, there is weak evidence for population differentiation in pulse amplitude – which showed little correlation with the slow/fast pulse traits (Figure S1). The pulse amplitude difference was not significant after applying permutation-based multiple test correction (indicating a 72% chance of getting at least one raw p-value as low as 0.088; Table S5). Figure 2 demonstrates that both of these traits had somewhat overlapping interquartile ranges, with more distinct distributions for pulse mode.

The above findings suggested that throughout the course of multiple courtship bouts in an uninterrupted mating setting, males from France strains were relatively more likely to utilize a fast pulse mode over a slow pulse mode. This trait’s population differentiation was also quantified by the Q_ST_ statistic, which yielded an elevated value of 0.797, which exceeds the F_ST_ values of 96.44% of examined SNPs genome-wide (see Materials and Methods). In general, Q_ST_ values that exceed the bulk of the genomic F_ST_ distribution are often considered potential targets of geographically variable directional selection, and our comparison conservatively used F_ST_ values from individual SNPs. Although this result should also be interpreted in the context of the number of traits initially examined, it suggests the possibility that the usage of fast versus slow pulses may have been influenced by differential selection between populations, whether acting on this or a pleiotropically correlated trait. Furthermore, the relatively consistent population differentiation observed for this trait makes it an appropriate target for QTL analyses involving phenotypically contrasting strains.

### SONG VARIATION BETWEEN RIL PARENTAL STRAINS REVEALS ADDITIONAL TRAITS FOR MAPPING

Next, we compared mean trait values between the two strains used in the establishment of our RIL panel, FR320N and ZI418N (“parental” strains) (Figure 2 – dotted rectangles; Table S5). Between these strains we found a total of 10 out of 37 traits with significant raw MW p-values (p<0.05) (Table S5). The two song traits mentioned above with regard to population-level differences, pulse mode preference and median pulse amplitude, were represented among this set of traits (Figure 2A & 2B, dotted rectangles). The similarity in magnitude and direction of parental differences in these traits has important consequences for our ability to interpret mapping results as representing likely contributors to population differentiation, particularly for the pulse mode trait.

Seven additional traits displayed quantitatively significant differences between parental strains: the number of bouts per minute, the transition probability of ‘null’ (no) song to sine song, the median sine and pulse train length, median sine amplitude, and mode carrier frequencies of sines and pulses (Table S4). Quantitative differences in these traits between parental strains imply that mappable variation could exist in the RIL panel, and we therefore elected to map these traits in addition to the two previously mentioned. While mapping efforts for most of these traits will not reveal loci that contribute to significant population differentiation between populations, they could reveal loci that make a meaningful biological contribution to the production of courtship behavior.

### SOME QTLS FOR TRAITS DIFFERENTIATED BETWEEN PARENTAL STRAINS INCLUDE KNOWN COURTSHIP GENES

We used a modified version of r/QTL (Broman *et al*. 2003; da Silva Ribeiro *et al*. 2024) to first map the song traits associated with parental strain but not population-level differences, under a single-locus QTL model. QTL mapping results for these traits are depicted in Figure 3 for the amount and type of song and in Figure 4 for the properties of pulse or sine song, with full LOD profiles given in Table S6. For two traits, mode sine carrier frequency and median sine amplitude, no genome-wide significant QTLs were obtained. Conversely, a total of 21 significant QTL peaks were identified for the remaining six traits, as described individually below (Table S7). Within each of these QTLs, we highlight below any known genes associated with male mating behavior (Table S2). However, not all QTLs include genes known to contribute to courtship behavior, and even where they are present it is possible that other genes with currently unknown roles in courtship behaviors may underlie some QTLs (Mackay 2009).

**Figure 3:**
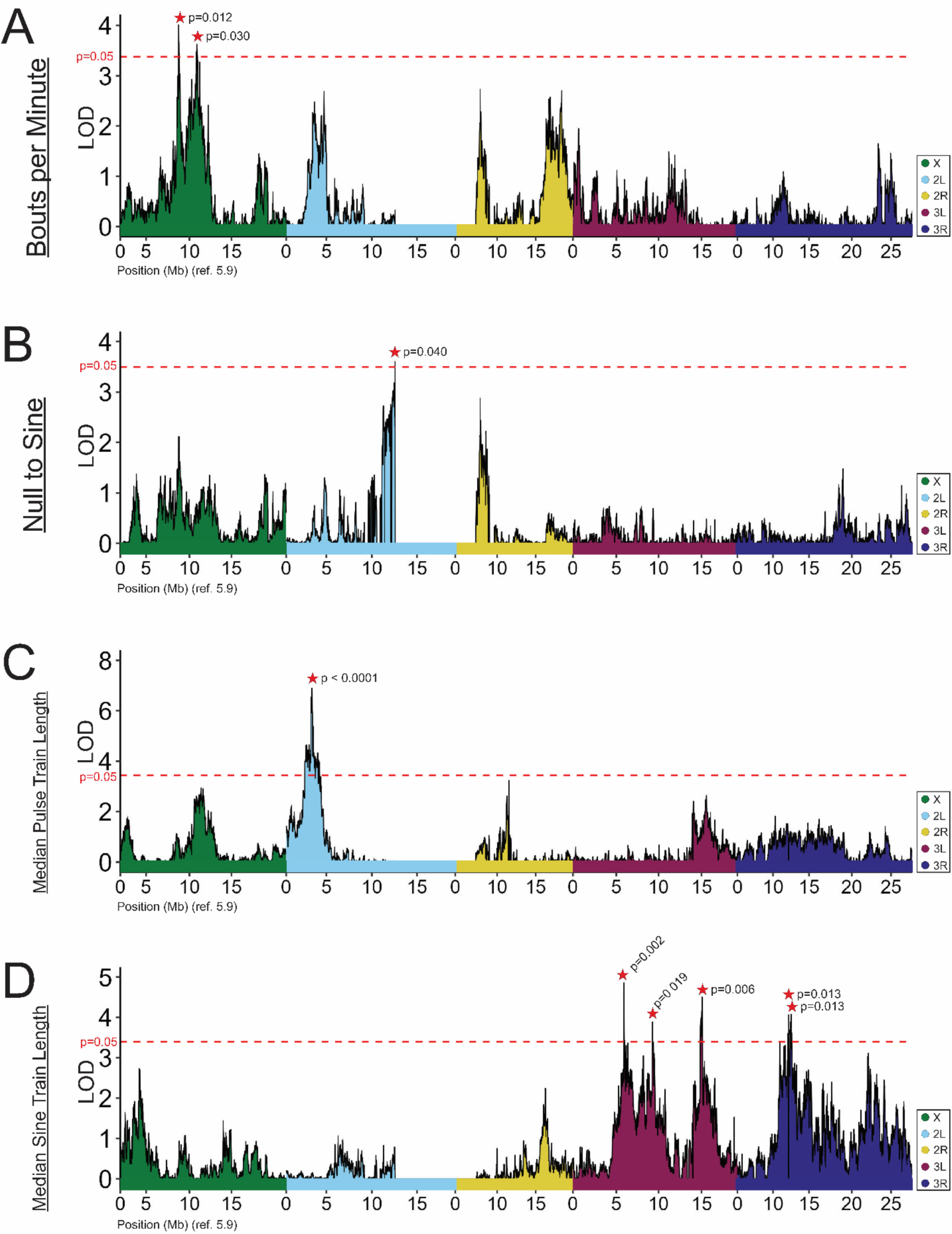
Traits associated with the amount and type of song map to unique QTLs. The number of bouts per minute (A) mapped to a pair of nearby QTLs on the X chromosome - the only X-linked QTLs detected in this study. The probability of initiating sine song (“Null to Sine”) (B) and the median length of pulse trains (C) mapped to single QTLs. In contrast, the median length of sine trains displayed a more polygenic basis, with multiple QTLs across both arms of chromosome 3, and distinct peaks from the pulse song counterpart of this trait (D). LOD values are plotted against *D. melanogaster* genome (release 5.9) positions, where color denotes chromosome arm. Genome-wide significance thresholds of LOD values corresponding to p = 0.05 for each scan are displayed as dashed lines. Significant peaks are denoted with a red star, with p-values noted. Further characteristics of each QTL are given in Table S7.

**Figure 4:**
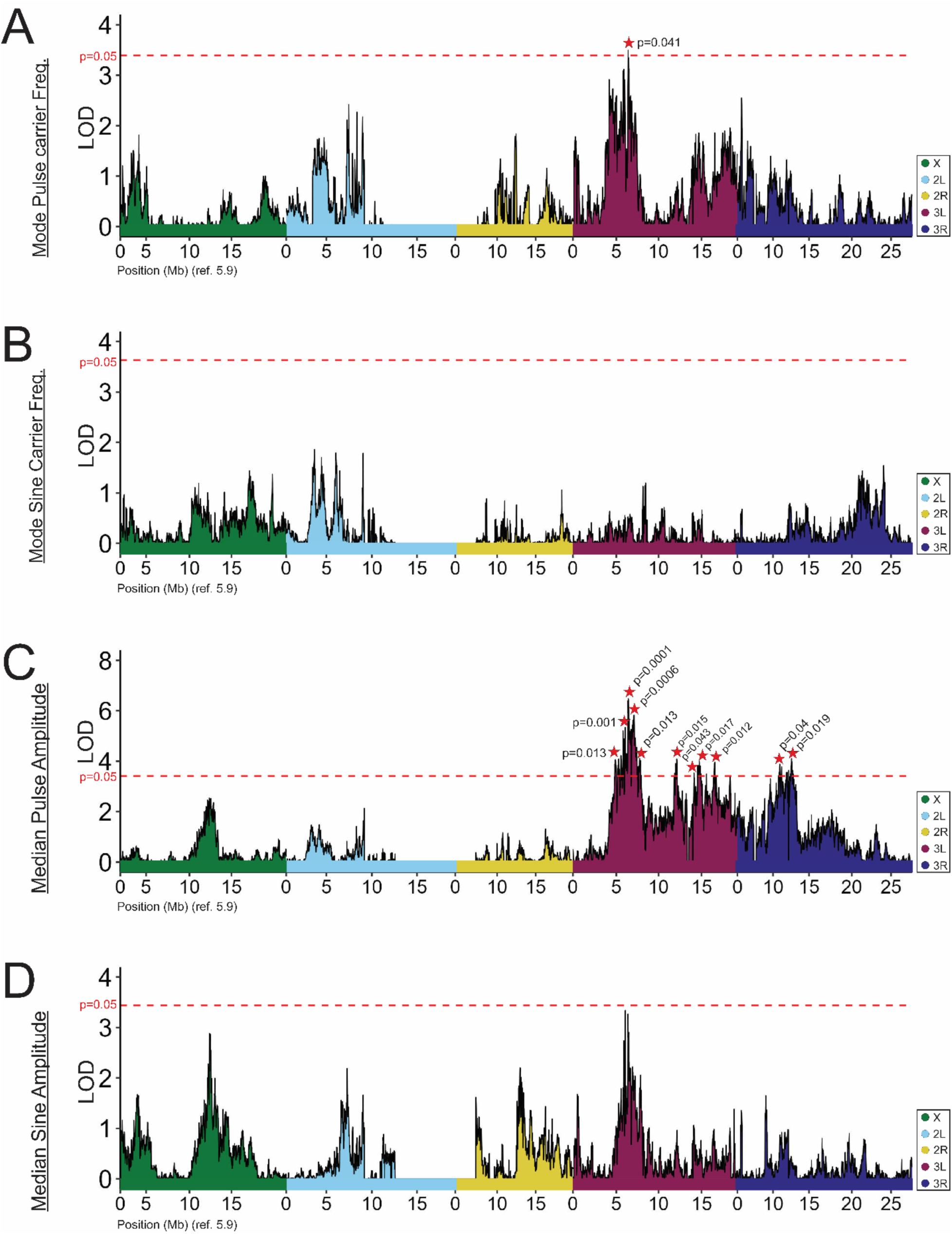
Traits associated with sine and pulse song qualities map to unique autosomal QTLs. Both pulse (A) and sine (B) song amplitude and carrier frequencies were distinct between RIL parental strains, and map to unique QTLs. The only instance of partially overlapping QTL peaks was a region on chromosome 3R, overlapping QTLs for pulse amplitude (C) and sine train length (D), while the QTL for pulse carrier frequency was nearby, as was the nearly significant peak for sine amplitude. LOD values are plotted against *D. melanogaster* genome (release 5.9) positions, where color denotes chromosome arm. Genome-wide significance thresholds of LOD values corresponding to p=0.05 are displayed as dashed lines. Significant peaks are denoted with a red star, with p-values noted. Further characteristics of each QTL are given in Table S7.

The average number of bouts per minute differed between the RIL parental strains (Table S5) [FR320N (6.24+/-2.832) vs. ZI418N (7.831+/-2.294)] and mapped to two primary QTLs on the X chromosome (Figure 3A; Table S7). The second interval (ChrX:10.6-11.4 Mb; note that all positions given here refer to *D. melanogaster* reference release 5.9) was found to include two courtship genes from our annotated list, *discs large 1* (*dlg1*, FBgn0001624) and *gustatory receptor 10b* (*Gr10b*, FBgn0030297).

Intuitively, a second parental-differentiated trait, the transition probability from null sound to sine song was found to be correlated with the above-discussed bouts per minute trait. As expected, this “Null to Sine” trait value was higher in the parental strain with greater bouts per minute [ZI418N (0.005+/-0.004) vs. FR320N (0.003+/-0.002)] (Figure 2; Table S5). In spite of the above-mentioned correlation with bouts per minute, the identified QTLs for these traits were distinct, although some non-significant peaks coincided (Figure 3B). The null to sine probability mapped to a region near the centromere on chromosome arm 2L starting at ∼12.1 Mb (Figure 3B). This QTL’s CI spans the low recombination centromeric region and extends until chromosome arm 2R at ∼7.04 Mb. Ten courtship-associated genes were found in this interval, 4 of which encode gustatory receptors (*Gr33a*, *Gr39a*, *Gr43a*, and *Gr47a*) (Table S8). Three genes that have been previously established to play roles in male courtship initiation were also found in this interval (Table S8). The gene *Chemosensory protein B 42a* (*CheB42a,* FBgn0053348) encodes a pheromone-binding protein that plays a role in the rate and speed of courtship initiation in males (Park *et al*. 2016). A neighboring gene, *pickpocket 25* (*ppk25*, FBgn0053349), was similarly implicated in the initiation of courtship (Liu *et al*. 2012). Although the simplest hypothesis would be that only a single gene is responsible for this QTL, the presence of genes relevant to courtship initiation is notable with regard to the transition probability to sine song.

The median lengths of sine and pulse trains were both significantly different between parental strains, and shared positive correlations with one another (Figure 1A & 1B). In both cases, train lengths were longer in the Zambian [sine: 0.094+/-0.032, pulse: 1.158+/-0.136] cohort and shorter in the French cohort [sine: 0.078+/-0.025, pulse: 1.069+/-0.128] (Figure 2; Table S5). Median pulse train length mapped to a single QTL with restrictive peak Cis (Figure 3C; Table S7) between Chr2L:3.21-3.39 Mb that does not harbor annotated male courtship-related genes. Whereas, we found that median sine train length mapped to several separate QTLs on chromosome 3: three peaks on the left arm and two peaks on the right arm (Figure 3D) (Table S7). A single QTL at Chr3R:12.1-12.6 Mb overlaps a known courtship gene, *spineless* (*ss*, FBgn0003513) (Table S8).

We also investigated trait variation in the mode carrier frequency of pulse and sine songs, which were higher in the France strain [211.7+/-40.7 versus 199.3+10.9 for pulses, 297.6+/-11.3 versus 291.9+/-5.5 for sines] (Figure 2; Table S5). Pulse carrier frequency mapped to a single primary QTL on chromosome 3L (Figure 4A & 4B; Table S7) and was not found to harbor known courtship genes. There was no significant QTL associated with sine carrier frequency, and underlying LOD landscapes suggest that, as with train lengths, sine and pulse carrier frequencies may be governed by unique loci (Figure 4A & 4B, Table S7). This trend is somewhat recapitulated in mapping the amplitude of pulse (Figure 4C) and sine song (Figure 4D), although no QTLs were detected for the latter trait.

Relative to the above traits, the median amplitude of pulses gave a relatively more polygenic signal, mapping to 11 unique QTLs that form roughly three clusters across chromosome 3 (Figure 4C; Table S7). The two most statistically significant peaks mapped to chromosome 3L (inclusive Cis 3L:5.98-6.71 Mb and 3L:6.80-7.24 Mb) and were flanked by two ‘minor’ peaks (Figure 4A; Figure S5; Figure S6; Table S7). The more significant of these two (peak 3) contains the courtship-associated gene *pale* (*ple,* FBgn0005626). *Pale* encodes a rate-limiting enzyme involved in catecholamine synthesis, such as the production of dopamine precursors, and has been demonstrated to play roles in a variety of behaviors, including courtship (Liu *et al*. 2009).

A second cluster on chromosome 3L harbored 5 additional QTLs, ranging between 3L:11 Mb and 3L:16.5 Mb and spaced roughly 1 Mb apart (Figure 4C). Peak 5 (Figure S6, Table S8) contains a second identified courtship-associated gene, the gustatory receptor *Gr68a* (FBgn0041231). The third cluster on chromosome 3R contained the final two QTLs, which each have confidence intervals within the 3R:10.5-13.4 Mb region. Because this trait showed possible evidence of population differentiation, for completeness we report a scan for local adaptation statistic outliers (Materials and Methods; Figures S5 & S6; Table S8; Table S9). However, we emphasize that the evolutionary forces underlying variation in this trait remain uncertain.

### COMPOSITE TRAIT ANALYSIS YIELDS UNIQUE QTL LANDSCAPES OF TWO SONG TRAITS

The extent of correlation among song traits (Figure 1) and occasional similarity in LOD landscapes of QTL maps suggested that some overlapping QTLs between traits could reflect pleiotropy. While full multivariate trait mapping departs from the focus of the present study on connections between genomic regions and specific aspects of song, we did investigate the potential for pleiotropy contributing to overlapping QTLs by performing residual trait mapping between specific pairs of traits. In general, for two traits with higher and lower LD scores involving their overlapping QTL peaks, we obtained residuals for the lower-LOD trait after performing a linear regression of its trait values against the higher-LOD trait. These residuals, which reflect the RIL variation in the lower trait not explained by its correlation with the higher trait, were then used in our standard QTL mapping pipeline.

One pair of positively correlated traits – median sine train length and median pulse amplitude (Figure 1) – had two overlapping peaks. Between these two pairs of overlapping QTLs, each trait had the higher LOD score in one case (Figure S2A; Figure S3; Table S10), and therefore regression and residual mapping were performed in both directions. When the residual trait values for median pulse amplitude were mapped, six chromosome 3 QTLs remained, one of which overlapped the original peak of interest, while another contained the courtship-associated gene *doublesex* (Table S9). When the residual trait values of median sine train length were mapped instead, all five of that trait’s originally significant chromosome 3 QTLs were eliminated, but a novel QTL peak on the X chromosome was detected [C.I. approx. X:4.14Mb-4.41Mb], which did not harbor any known annotated courtship genes (Figure S3A). Hence, among the two intially overlapping peaks, there is evidence for a locus underlying pulse amplitude that can not be explained by that trait’s overlap with sine train length, whereas the QTL signal observed for sine train length could be explained via pleiotropy with pulse amplitude.

Another instance of QTL peak overlap occurred between the pulse carrier frequency and median pulse amplitude traits, which shared an overlap within the pulse carrier frequency single significant peak (Figure 4A, 4C; Table S7). When fit to median pulse amplitude, the residual trait values of pulse carrier frequency did not yield significant QTL (Figure S3C). This result may imply that the QTL signal for pulse carrier frequency is driven primarily by its pleiotropic relationship with median pulse amplitude. Our results are consistent with previous evidence of a positive relationship between these traits (Clemens *et al*. 2018), and that males with silenced motoneurons produce pulse song that is decreased in both amplitude and carrier frequency (Shirangi *et al*. 2014).

Lastly, we performed a linear regression between median sine amplitude and median pulse amplitude song traits (Figure S4). These traits displayed the strongest positive correlations in both inbred and RIL datasets (Figures 1, S1, S2). These traits did not share overlapping QTLs (initial mapping of median sine amplitude did not reveal any significant QTL). However, they did share very similar overall QTL landscapes, and a nearly-significant QTL for sine amplitude aligned closely with the strongest QTL for pulse amplitude (Figures 4C and 4D). Residual mapping revealed a novel QTL for median sine amplitude on chromosome 3R [C.I. approx. 3R:19.35Mb-19.98Mb], which does not harbor known annotated courtship genes. (Figure S4). When comparing the initial sine amplitude QTL landscape to the residual trait landscape, we found that many of the non-significant peak signals on chromosome 3L were “flattened” or had diminished signal in the residual map, consistent with much of the trait signal on chromosome 3L being driven by one or more genetic factors underlying the correlation of these two amplitude traits.

### A SONG TRAIT DIFFERENTIATED BETWEEN FRANCE AND ZAMBIA POPULATIONS YIELDS A SINGLE STRONG QTL

We next mapped the song trait that represented the most geographically differentiated trait among those studied (Figure 2; Table S5). The ratio of slow pulses to the total number of pulses, which was significantly higher in the Zambian strains relative to France strains (0.833+/-0.067 vs. 0.725+/-0.125, respectively) (Figure 2A; Table S5). One major QTL peak on chromosome 3R was identified with a broad Cl interval at approximately 3R:20.0..22.6 Mb (Figure 5A; Table S7), which was estimated to explain 42% of the parental strain trait difference.

**Figure 5:**
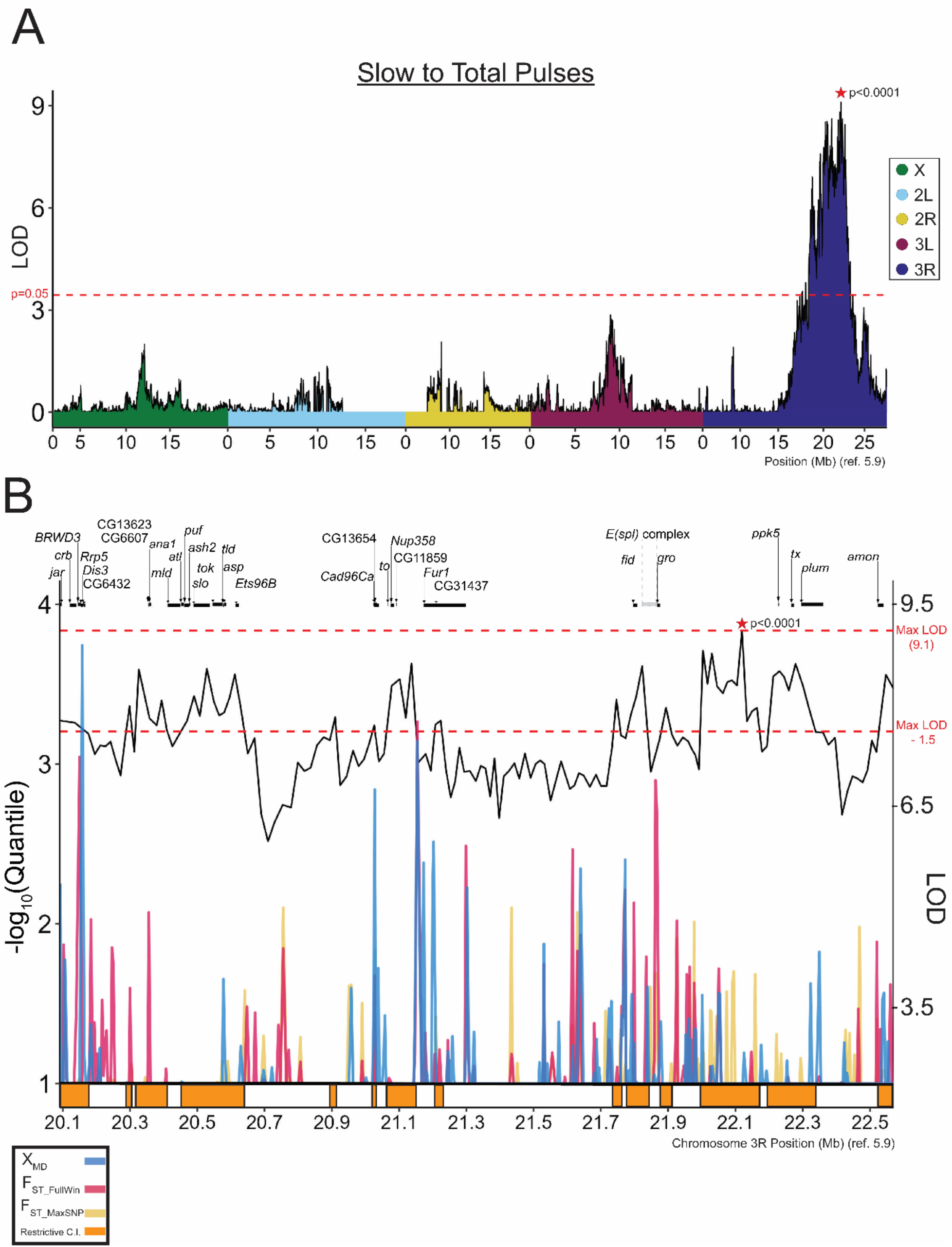
Pulse mode preference maps to a single strong effect QTL, with potentially relevant genes indicated by functional annotation and population genetic analysis. (A) QTL mapping of the quantitative trait “slow to total pulses” in song reveals a single strong QTL on chromosome 3R with a complex LOD landscape. All features of this plot are as described for Figures 3 and 4. (B) Negative log quantile values for *F*_ST_FullWin_ (Red), *F*_ST_MaxSNP_ (yellow), and *χ*_MD_ (blue) reveal genomic regions with elevated population differentiation quantile values, sometimes overlapping peak LOD windows. -log_10_ quantile values on the left *y*-axis begin at 1.00 to better visualize outlier windows. Window LOD scores are represented as a black line (right *y*-axis). Restrictive confidence intervals are represented as orange boxes under the *x*-axis. The QTL peak’s maximum LOD score and the (MaxLOD – 1.5) threshold are displayed as dashed red lines across the y-axis, with the latter value representing a threshold for restrictive Ci inclusion. We find an aggregation of more differentiated genomic sites within restrictive QTLs around 3R:21.75-22.25 Mb. This genomic region has been previously implicated in courtship behavior (Moehring & Mackay 2004). Further gene annotation details are available in Table S8.

This trait was estimated to have approximately a 90% probability of reflecting genuine population differentiation, and presented a *Q_ST_* value consistent with the differential action of directional selection (on either this or a pleiotropically correlated trait) between populations. Under the hypothesis that a gene underlying a trait with elevated population differentiation might display elevated genetic differentiation, we chose to supplement our QTL mapping with a population genetic analysis of the detected QTL region, in which we identified genomic windows within the QTL’s restrictive CIs that were associated with values of either window *F_ST_*, maximum SNP *F_ST_* (da Silva Ribeiro *et al*. 2022) or the Comparative Haplotype Identity statistic (Lange & Pool 2016) within the top 1% on the chromosome arm (Materials and Methods; Table S9). We also made note of an expanded range of candidate genes on functional grounds, including nervous system genes and those related to wing development.

The highest LOD scores within this QTL were amongst a group of restrictive CI regions (*i.e.* intervals within 1.5 LODs of the QTL maximum) toward the distal/right side of this QTL. Within one of these restrictive CI regions, we identified a cluster of outlier statistics around 3R:21.75-22 Mb (Figure 5B, orange intervals; Materials and Methods; Table S7). Within this gene-dense region is the *enhancer of split* complex and flanking genes, which have been previously implicated in the quantitative trait mapping of courtship behavior (Moehring & Mackay 2004). Additionally, the transcriptional corepressor *groucho* (*gro*, FBgn0001139) falls within this region (Figure 5B), and has been implicated in the *retained*-mediated repression of transcriptional targets involved in male courtship behavior (Shirangi *et al*. 2006).

Two additional male courtship genes were found within other restrictive CI regions of this same QTL (Figure 5B; Table S8). Near the middle of the full QTL region, the gene *takeout*, a lipid binding protein that is a target of the sex determination pathway and implicated in male courtship (Dauwalder *et al*. 2002), is flanked by highly differentiated windows (Figure 5B). Toward the proximal/left side of the QTL, the *slowpoke* gene, which has a demonstrated role in sine song carrier frequency variation in the *D. simulans* clade (Ding *et al*. 2016) is present within the restrictive CI. This gene does not overlap or adjoin any population genetic outlier windows (Figure 5B), but this pattern does not exclude the possibilities of elevated differentiation at more distant regulatory regions or a causative locus that lacks a strong population genetic signal.

Three additional regions of this QTL’s restrictive CIs were found to contain population genetic outlier windows (Figure 5; Table S9). The first window contains the gene *jaguar* (*jar*, FBgn0011225). The second window overlaps three genes: *BRWD3*, *Rrp5*, and *Dis3* (FBgn0011785, FBgn0039182, FBgn00339183). The third window does not contain any genes, but sits upstream of the first exon of the closest proximal gene, *Furin-1* (*Fur1,* FBgn0004509). This gene has known roles in synaptic transmission at neuromuscular junctions (Kim *et al*. 2015), and its disruption via RNA-interference is associated with abnormal locomotion and flight (Schnorrer *et al*. 2010). Although we highlight multiple potential genes that may underlie this QTL, we note that it may be driven by any one of them, by more than one of them, or by none of those listed above.

## DISCUSSION

Here we describe natural variation in male courtship song components between two natural populations of *D. melanogaster* from contrasting environments. Comparison of population cohorts revealed significant phenotypic differences in pulse mode preference, and relatedly the total number of fast pulses performed in the assay. We found that the slow to fast preference is partially explained by a QTL with moderate effect size. Conversely, the median amplitude of pulses, which had limited support for population differentiation, was found to be under the control of multiple QTL intervals of relatively small effects. Other traits differentiated between the RIL parental strains had anywhere from zero to six significant QTLs detected. However, in light of the total effect sizes estimated across QTLs for each trait (Table S7), along with the finite statistical power expected for detecting QTLs from a few hundred RILs (*e.g.* King *et al*. 2012), we would predict that each of these traits is also influenced by an unknown number of smaller effect loci that did not achieve genome-wide significance in our mapping study.

### Phenotypic variability among RIL parental strains reveals unique QTL for multiple courtship traits

Mapped traits that did not have significant differences at the population level but did display significant parental strain differences were mapped in an attempt to identify genes that may regulate courtship behavior. Components of courtship behavior are responsive to artificial selection in lab settings using wild-derived populations (Ritchie & Kyriacou 1994; Gaertner *et al*. 2015), reinforcing the view that fitness-relevant variation in courtship behaviors may be present within natural populations. The generation of a RIL panel from two inbred strains will capture a limited quantity of naturally segregating variation, but the origins of our parental strain from two distinct natural populations may have enhanced the extent of trait variation in behavior that is amenable to QTL mapping.

Aside from pulse mode preference, we found that four of eight parental-differentiated traits yielded significant QTLs overlapping known courtship-related genes (Table S2; Table S8), and our results provide a preliminary association of these male song components with previously implicated genes. Most of the significant QTL were found on autosomes (Figures 3 & 4), with only two neighboring QTLs for the same trait identified on the X chromosome. (Y chromosome effects were not considered in the present study, due to the absence of available RIL sequencing data from males) While the total number of QTLs and the linkage that exists among some of them preclude strong conclusions regarding X-linkage, a reduced occurrence of X-linked QTLs might be predicted if genes involved in male song production are sometimes subject to sexually antagonistic selective pressures (Snook *et al*. 2005), in light of the X chromosome spending two thirds of its time in females.

In general, we found that even when corresponding pulse and sine song traits both differed between RIL parental strains, they mapped to distinct genomic intervals. This pattern held for pulse versus sine train length (Figure 3C & 3D; Table S7) and for pulse versus sine carrier frequency (Figure 4A & 4B; Table S7), in agreement with the lack of strong correlations between these traits (Figure S1). This evidence for distinct genetic architectures, in spite of co-directional differences in these pulse and sine song traits between our France and Zambia parental strains, supports previous findings that components of sine and pulse song can have unique genetic (and neuronal) bases (Shirangi *et al*. 2013; Ding *et al*. 2016; Hiroshi *et al*. 2022; Shiozaki *et al*. 2023; Lillvis *et al*. 2024). In contrast, we found that the nearly genome-wide significant (p = 0.06) peak for sine amplitude (Figure 4D) falls within the inclusive CI of the most significant QTL for pulse amplitude, at roughly 3L:6.30 Mb – 6.32 Mb (Figure 4C; Table S7, peak 3). The potentially overlapping architecture of the two amplitude traits may indicate a pleiotropic basis and aligns with the stronger correlation between these traits within both populations (Figure S5, S6).

### The rate of slow to fast pulses is the most phenotypically differentiated song trait between Zambia and France populations and involves a single detectable QTL

Among the 37 song traits examined here, two traits related to pulse mode usage, particularly the preference for slow versus fast pulses, received the strongest statistical support for differentiation between the France and Zambia populations examined. The distinction between fast and slow pulse modes is a relatively recent discovery and its possible role in sexual selection is currently unknown, although the trait does seem to affect female movement responses to song (Clemens *et al*. 2018). This same study showed that males use sensory input (primarily visual) to bias which pulse mode is used (Clemens *et al*. 2018).

Assay conditions, such as chamber size and female behavioral plasticity are thus an important consideration when interpreting results regarding this trait. Recording chambers used in the study were small (Arthur *et al*. 2013), constraining flies to a minimal space. We did not track the positions of flies during courtship, so we cannot rule out that male song varied in response to female signals. The difference in pulse mode preference in the present study may either represent a cumulative difference in male response to females, or simply represent a difference in a probabilistic preference of pulse choice when other environmental parameters are held constant. Still, QTL signals for this trait were very strong (Figure 5A) and suggest that population genetic differentiation underlies trait differences.

Our analysis provides evidence that pulse usage may have been influenced either directly or indirectly by population-specific directional selection. In a conservatively designed comparison between *Q_ST_* and *F_ST_*, the pulse type ratio showed greater differentiation than all but around 3.5% of genome-wide SNPs. If pulse type usage has in fact been influenced by recent directional selection, the reasons remain unclear. Although female preference varies between some southern African and European strains, there is no documented role of courtship song in M/Z-type mating preference (Colegrave *et al*. 1999). Hence, it is unclear to what degree any directional selection has resulted in song differentiation that impacts male reproductive fitness. However, it is worth emphasizing that all experiments in this study were conducted at room temperature. In contrast, the far more variable temperatures encountered by natural populations, especially in the temperate zone, may exert plastic influences on song components (Stanley & Kyriacou 2021), and it is possible that genetic adaptation may allow for the production of universally favorable song characteristics across divergent environments, even if some specific song differences persist at a given temperature.

Alternatively, it is possible that directional selection, if present here, may not have been acting upon song at all. Population genomic scans for selection in this species have reported evidence for local adaptation between warm- and cold-adapted natural populations for multiple categories of genes that modulate synaptic function, including neuropeptide receptors, ion channels, and structural components of the synapse (*e.g.* Pool *et al*. 2017). It is possible that the adaptive evolution of one or more aspects of synaptic function, even if driven by selection to maintain nervous system function in a novel environment, might have pleiotropic consequences for behaviors such as those involved in courtship song. Adaptive responses to other selective pressures, such as insecticides, some of which target insect nervous systems, also may trigger pleiotropic effects on traits that are not the direct target of selection. Hence, further study will be needed to assess whether pulse mode usage has been targeted or pleiotropically influenced by directional selection between African and European *D. melanogaster* populations.

The pleiotropy hypothesis could potentially also explain variation in amplitude (*i.e.* volume), which showed weaker evidence of differentiation between populations. Pulse amplitude was found to be somewhat higher in France than Zambia males (Figure 2; Table S5), sine amplitude showed a lesser trend in the same direction, and both traits were significantly higher in the France parental strain than in the Zambia parent (Table S5). Flies from these populations tend to be slightly larger (Lack *et al*. 2016b). Based on measurements of females from that study, the France RIL parent has 8.1% greater thorax length and 5.5% greater wing width than the Zambia RIL parent. If larger flies tend to sing with greater volume, then differences in amplitude could be downstream of differences in body or wing size. Empirically, this relationship is unclear, although there is indirect evidence that physical size of wing components, such as the musculature, can play roles in the sexually dimorphic expression of male song (Shirangi *et al*. 2013). Further research will be needed to determine the degree to which song variation and differentiation within *D. melanogaster* is shaped by directional selection to modify or maintain sexually-selected traits in populations experiencing different environments, directional selection on pleiotropically correlated traits, stabilizing selection, genetic drift, or other forces.

We identified a single large-effect QTL for pulse mode usage on chromosome arm 3R. Curiously, this QTL for the trait with the strongest evidence for population differentiation had a stronger mapping signal than any of the 21 QTLs identified for any of the other traits that differed between the RIL parent strains. This QTL had a somewhat bimodal shape (Figure 5A) that could reflect either stochastic factors in QTL mapping or else the contribution of more than one locus.

In *Drosophila*, an average QTL may encompass hundreds of genes (Mackay 2001). Functional approaches, such as complementation tests or gene disruption experiments, have traditionally been employed to identify candidate genes from QTL intervals, but can become a laborious task when attempting to test all possible candidates within a QTL CI. We thus supplemented our QTL mapping with a population genetic analysis of QTL intervals, leveraging available population genomic data (Lack *et al*. 2016a) to identify specific loci with elevated genetic differentiation between populations that might contribute to elevated trait differentiation. The candidates identified for pulse model preference, such as the region surrounding the *enhancer of split* complex, *groucho*, *takeout*, and other genes (Figure 5B; Table S8), provide promising candidates to be tested in future studies. Subsequent analyses such as fine mapping and functional testing such as via reciprocal hemizygosity tests (Stern 2014) may ultimately confirm specific genes that contribute to population differences in courtship song.

Collectively, our results provide evidence of variation in pulse song usage between naturally occurring populations of *D. melanogaster*. By combining RIL-based QTL mapping with population genetic inference, we identified a set of genes that may be tested in future studies. We also note that all 37 traits examined showed meaningful variation among the studied strains, further reinforcing the vast potential of natural variation for investigating the genetic mechanisms underlying behavior.

**Table 1:** Summarized Results for Mapped Traits. Outcomes of population comparisons, RIL parental strain comparisons, and QTL mapping are listed for one trait with clear evidence of population differentiation (Slow to Total Pulses, bottom row) and eight other traits differing between RIL parental strains. FR and ZI indicate the France and Zambia population samples, from which multiple inbred strains were studied. “MW-P” indicates Mann-Whitney U Test p-values, presented for both the population and parental strain comparisons. The range of QTL effect size point estimates (proportion of parental strain difference explained) among each trait’s significant QTLs is presented; however, we note that some estimates for multi-QTL traits are likely to be inflated by linkage to other QTLs. Candidate genes based on functional considerations (see Materials and Methods) are listed for each trait. Genes listed for the trait “Slow to total pulses” additionally include others that fell within population genetic statistic outlier windows (Table S8, S9; Materials and Methods). More detailed results, including additional gene annotations, are provided in Tables S7, S8, and S9.

| Trait | FR Mean | ZI Mean | Pop. M-W P | FR Parent | ZI Parent | Par. M-W P | # QTLs | Effect range | Candidate genes |
| --- | --- | --- | --- | --- | --- | --- | --- | --- | --- |
| Bouts per Minute | 7.00 | 7.05 | 1 | 6.24 | 7.83 | 0.0077 | 2 | 0.393 - 0.431 | *dlg1, Gr10b* |
| Median Pulse Amplitude | 0.100 | 0.078 | 0.088 | 0.097 | 0.063 | 7.00E-06 | 11 | 0.266 - 0.402 | *Abd-B, Gr68a, mir-274, mir-285, mir-956, ple* |
| Median Pulse Train Length | 1.16 | 1.09 | 0.161 | 1.07 | 1.16 | 0.0054 | 1 | 0.863 | none |
| Median Sine Amplitude | 0.177 | 0.171 | 0.600 | 0.19 | 0.14 | 0.0007 | 0 | n/a | n/a |
| Median Sine Train Length | 0.806 | 0.892 | 0.133 | 0.78 | 0.94 | 0.0313 | 5 | 0.499 - 0.747 | *ss* |
| Mode Pulse Carrier Freq. | 200 | 203 | 0.230 | 212 | 199 | 0.0140 | 1 | 0.270 | none |
| Mode Sine Carrier Freq. | 289 | 291 | 0.887 | 298 | 292 | 0.0026 | 0 | n/a | n/a |
| Null to Sine Trans. Prob. | 0.0039 | 0.0040 | 0.962 | 0.003 | 0.005 | 0.0008 | 1 | 0.430 | *btv, CheB42a, Gr33a, Gr39a, Gr43a, Gr47a, jhamt, lig, Pal1, ppk25* |
| Slow to Total Pulses | 0.745 | 0.813 | 0.007 | 0.745 | 0.833 | 0.0001 | 1 | 0.423 | *ana1, BRWD3, Cad96Ca, CCAP-R, CG12290, CG13623, CG13654, CG31437, CG6432, CG6607, Dis3, [E(spl) gene cluster], fid, Fur1, gro, jar, mld, Nf1, ppk15, Rrp5, slo, to* |

## ACKNOWLEDGMENTS

We thank Ben Arthur and Joshua Lillvis for assistance with song analysis and members of the Pool Lab for helpful comments on this manuscript. This work was funded by NIH grant R35 GM13630 to J.E.P. and by the HHMI Janelia Research Campus Visitor Project Program.

## DATA AVAILABILITY STATEMENT

All raw trait data is provided in this article’s Supplemental Tables. No new genomic sequencing data was generated for this study. RIL sequence data reported by da Silva Ribeiro *et al*. (2024) is available from the NIH Sequence Read Archive (BioProject: PRJNA861300).

