## Supplementary material for "Courtship song differs between African and European populations of *Drosophila melanogaster* and involves a strong effect locus": Figure S

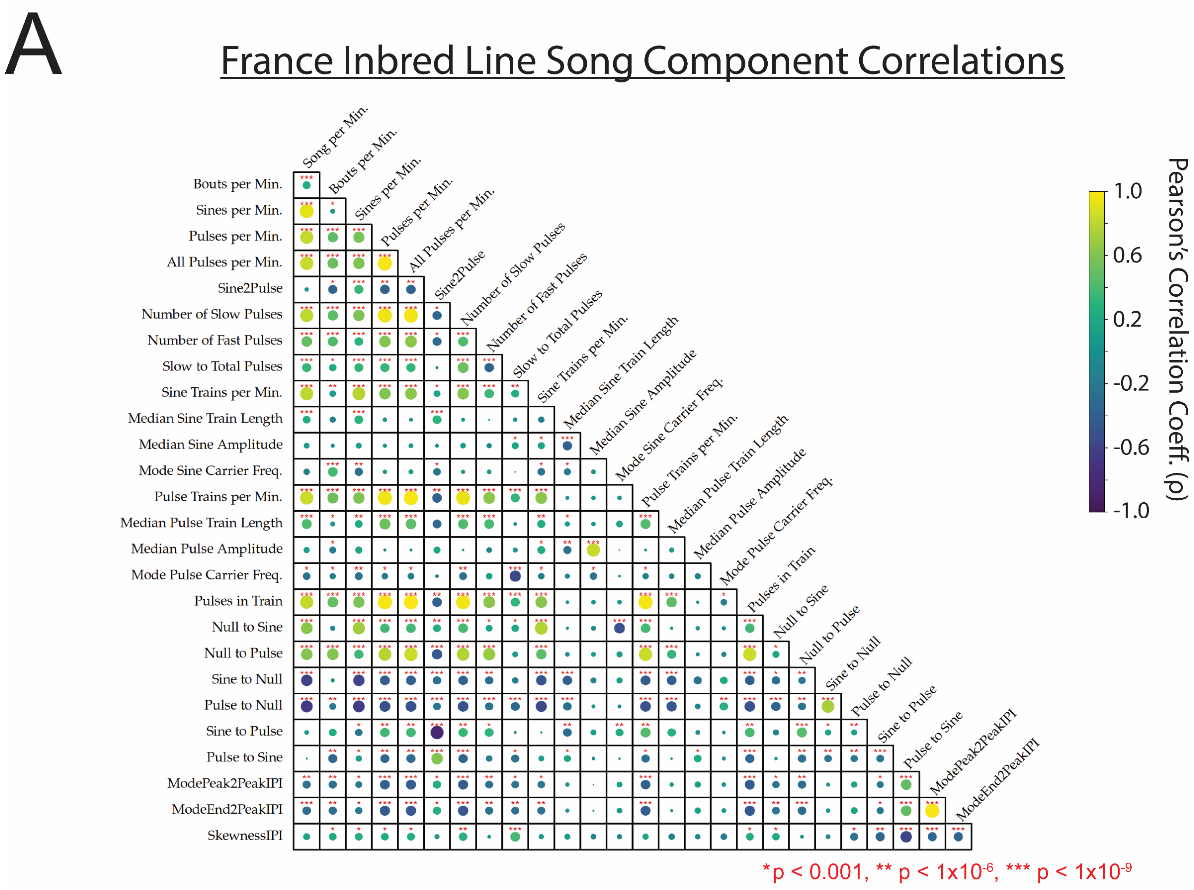
A
France Inbred Line Song Component Correlations
Song per Min.
Bouts per Min.
Sines per Min.
Pulses per Min.
All Pulses per Min.
Sine2Pulse
Number of Slow Pulses
Number of Fast Pulses
Slow to Total Pulses
Sine Trains per Min.
Median Sine Train Length
Median Sine Amplitude
Mode Sine Carrier Freq.
Pulse Trains per Min.
Median Pulse Train Length
Median Pulse Amplitude
Mode Pulse Carrier Freq.
Pulses in Train
Null to Sine
Null to Pulse
Sine to Null
Pulse to Null
Sine to Pulse
Pulse to Sine
ModePeak2PeakIPI
ModeEnd2PeakIPI
SkewnessIPI
1.0
0.6
0.2
-0.2
-0.6
-1.0
Pearson's Correlation Coeff. (ρ)
*p < 0.001, ** p < 1x10-6, *** p < 1x10-9

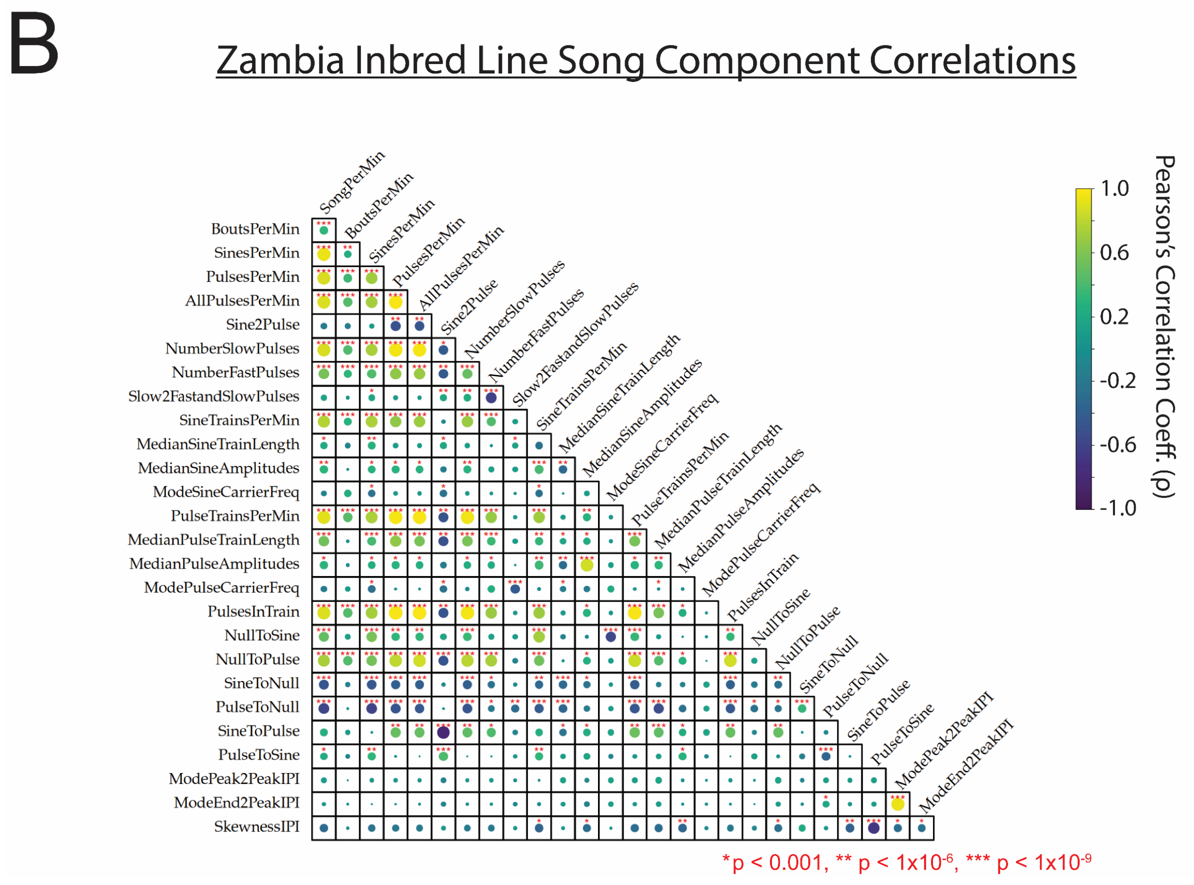
B
Zambia Inbred Line Song Component Correlations
SongPerMin
BoutsPerMin
SinesPerMin
PulsesPerMin
AllPulsesPerMin
Sine2Pulse
NumberSlowPulses
NumberFastPulses
Slow2FastandSlowPulses
SineTrainsPerMin
MedianSineTrainLength
MedianSineAmplitudes
ModeSineCarrierFreq
PulseTrainsPerMin
MedianPulseTrainLength
MedianPulseAmplitudes
ModePulseCarrierFreq
PulsesInTrain
NullToSine
NullToPulse
SineToNull
PulseToNull
SineToPulse
PulseToSine
ModePeak2PeakIPI
ModeEnd2PeakIPI
SkewnessIPI
1.0
0.6
0.2
-0.2
-0.6
-1.0
Pearson's Correlation Coeff. (ρ)
*p < 0.001, ** p < 1x10-6, *** p < 1x10-9

**Supplementary Figure 1: Correlations between traits scored in male song recordings from inbred strain data for population comparisons.** Correlation was assessed separately between France (**A**) and Zambia (**B**) sampling cohorts. Circle size indicates the relative strength of significance in Pearson's (*r*) covariance between traits, where the presence of a circle indicates minimally a p-value of $p < 0.05$, and correlations with an estimated multiple-testing corrected significant p-values of $p < 1.0 \times 10^{-3}$, $p < 1.0 \times 10^{-6}$ , and $p < 1.0 \times 10^{-9}$ are denoted with an asterisk(s) (*, **, and ***, respectively). Generally, significant correlations were among traits that had an *a priori* expectation of some correlation (*e.g.* the amount of song per minute positively correlating with the number of song bouts per minute).

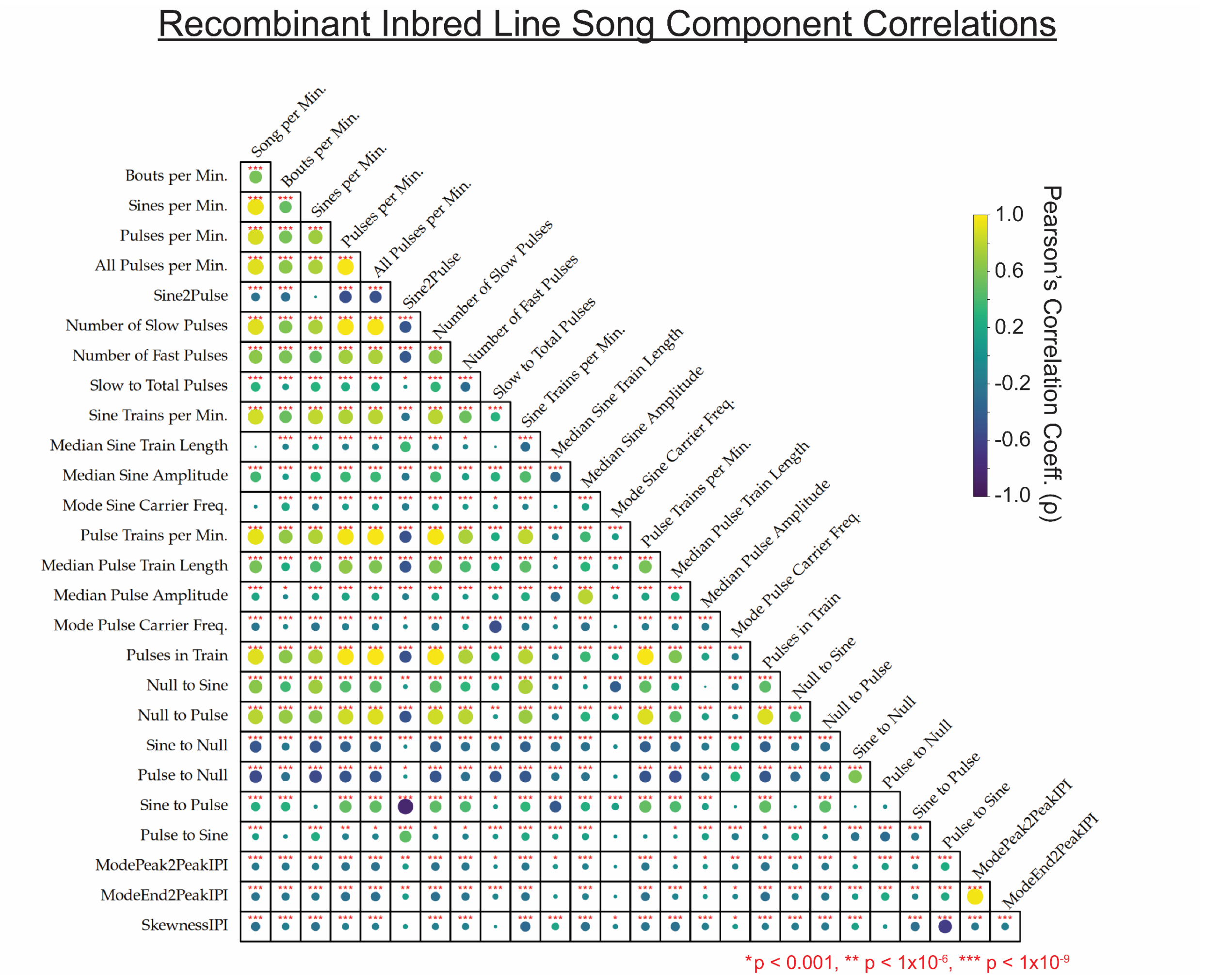


**Supplementary Figure 2: Correlations between traits scored in male song recordings from aggregate recombinant inbred line data.** Circle size indicates the relative strength of significance in Pearson's (*r*) covariance between traits, where the presence of a circle indicates minimally a p-value of $p < 0.05$, , and correlations with an estimated multiple-testing corrected significant p-values of $p < 1.0 \times 10^{-3}$, $p < 1.0 \times 10^{-6}$ , and $p < 1.0 \times 10^{-9}$ are denoted with an asterisk(s) (*, **, and ***, respectively).

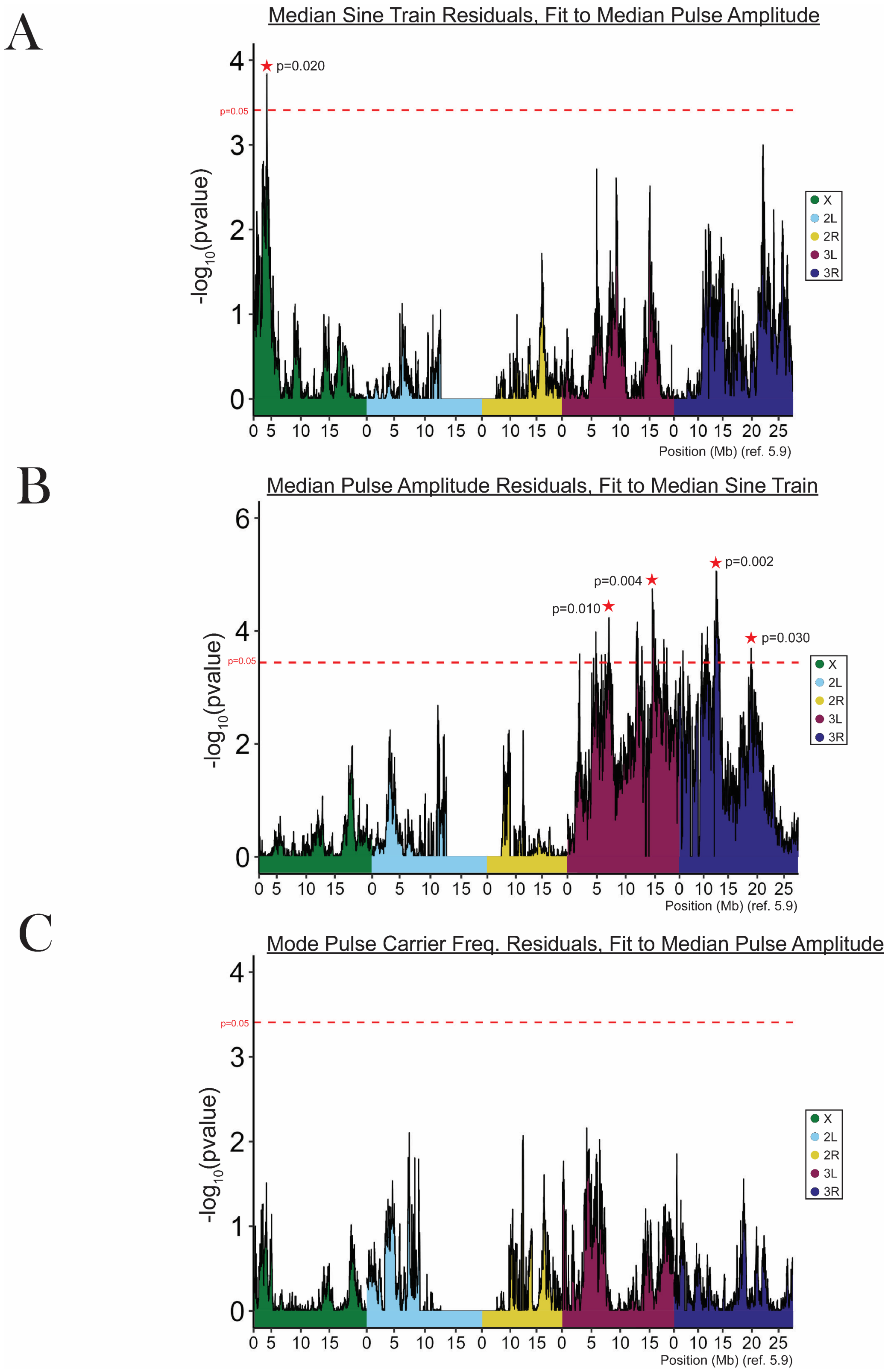


**Supplementary Figure 3: Residual trait QTL mapping suggests a pleiotropic basis of shared QTLs.** To assess whether overlapping QTLs from correlated traits might have a shared genetic basis, we performed a linear regression to obtain the residuals for the trait with the lesser maximum LOD score after accounting for its correlation with the trait having the larger LOD score among the overlapping peaks, and then running the residual values through our QTL mapping pipeline. (**A**) Median sine train length was fit to pulse amplitude overlap (Table S7, chromosome 3L peak around 15 Mb). The previously overlapping QTL and other QTLs along chromosome 3 (see Figure 3D) were no longer identified by residual trait mapping, suggesting the signals for these QTLs may have been largely due to factors also correlated with pulse amplitude. Rather, a peak on the X chromosome with some non-significant signal in previous mapping (Figure 3D) was identified as the single significant peak region associated with median sine train length residuals. (**B**) In the opposite analysis, pulse amplitude was fit to median sine train length, in order to focus on a second overlapping peak, near 13 Mb on chromosome 3R (Table S7). Here, some significant chromosome 3 peaks are present, but the initial overlapping peak is not. Instead, a novel peak on chromosome 3L harbored the only courtship-associated gene *doublesex* (Table S10), which was not noted in previous mapping (Figure 3C; Table S7). (**C**) Mode pulse carrier frequency was fit to median pulse amplitude. Here, a previously overlapping QTL on chromosome 3L (Figure 3A, 3C) was no longer detected, and residual mapping did not reveal any newly significant QTLs.

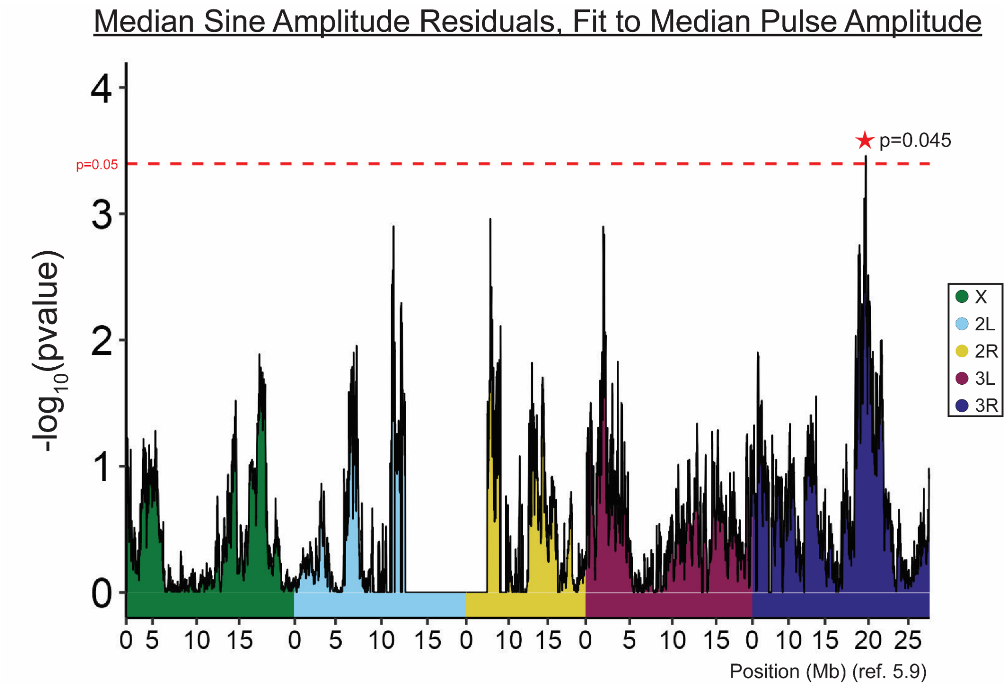


**Supplementary Figure 4: Residual trait QTL mapping reveals novel peak between highly correlated traits, median sine and pulse amplitudes.** Sine and pulse amplitude traits were the among the most positively correlated traits mapped in the present study, in both inbred and RIL datasets (Figures 1, S1, S2). Due to high correlation, similarity in traits, and similar QTL mapping landscapes (Figures 4C and 4D), we also attempted to map the residuals of sine amplitude fit to pulse amplitude. We found that an overlapping nearly-significant peak on chromosome 3L (Figure 4D) was absent from residual mapping results, suggesting this signal may be influenced by factors correlated with pulse amplitude. A single, novel significant peak was revealed on chromosome 3R, which shares no overlap with previously mapped traits and contains no annotated courtship genes.

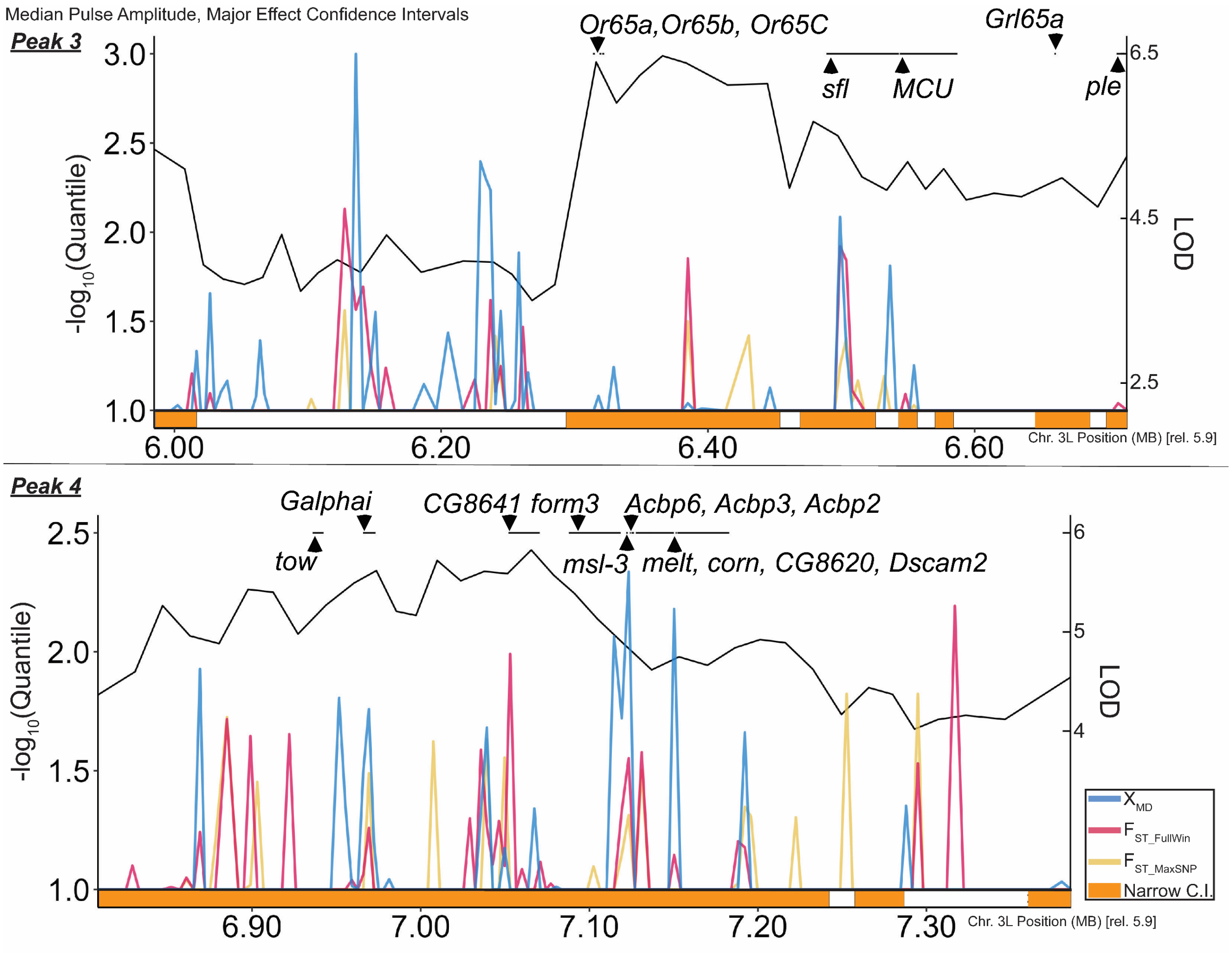


**Supplementary Figure 5: For two median pulse amplitude QTL intervals, the LOD landscape, population genetic outlier scan, and locations of selected genes are shown.** QTL mapping LOD scores are depicted as black lines, scaled according to the right-hand *y*-axis. Portions of the inclusive CI that qualified as being within the restrictive CI are shown as orange boxes along the *x*-axis, which is otherwise labeled based on megabase position in *D. melanogaster* genome release 5.9. Each of the three population genetic statistics used to quantify genetic differentiation between the France and Zambia population samples is shown in a distinct primary color (see legend at lower right). The quantile value of each analyzed window indicates the proportion of all windows on the same chromosome arm yielding a higher value for a given statistic. These quantiles are depicted according to a $-\log_{10}$ scale, such that lower quantiles (indicating more extreme high values for these statistics) correspond to peaks on this plot, and only quantiles less than 0.1 are visible (since the left *y*-axis starts at 1.0). We note that pulse amplitude showed possible evidence for population differentiation, but the possible role of adaptive evolution in shaping this trait's variation requires further investigation. Finally, the locations of genes corresponding to the criteria detailed in the Materials and Methods are shown along the top of each plot.

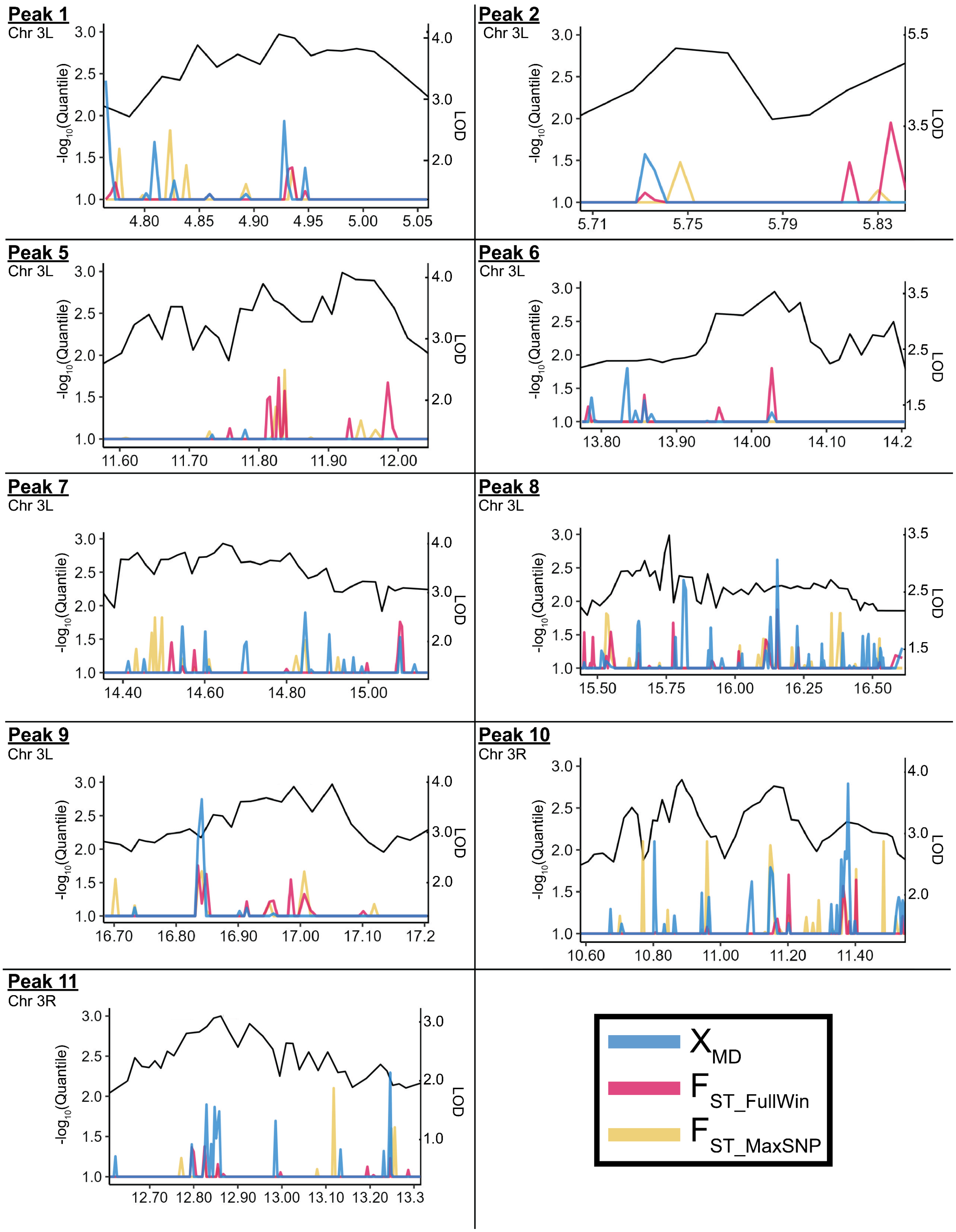
Peak 1
Chr 3L
-log10(Quantile)
LOD
4.80 4.85 4.90 4.95 5.00 5.05
Peak 2
Chr 3L
5.71 5.75 5.79 5.83
Peak 5
Chr 3L
11.60 11.70 11.80 11.90 12.00
Peak 6
Chr 3L
13.80 13.90 14.00 14.10 14.2
Peak 7
Chr 3L
14.40 14.60 14.80 15.00
Peak 8
Chr 3L
15.50 15.75 16.00 16.25 16.50
Peak 9
Chr 3L
16.70 16.80 16.90 17.00 17.10 17.2
Peak 10
Chr 3R
10.60 10.80 11.00 11.20 11.40
Peak 11
Chr 3R
12.70 12.80 12.90 13.00 13.10 13.20 13.3
X_MD
F_ST_FullWin
F_ST_MaxSNP

**Supplementary Figure 6: For nine median pulse amplitude QTL intervals, the LOD landscape, and population genetic outlier scan are shown.** All quantities are depicted in the same manner described in the legend of Supplementary Figure 2 above. Gene annotations are available in Table S8.
